# Females translate male mRNA transferred during mating

**DOI:** 10.1101/2023.09.22.558997

**Authors:** Luciano M. Matzkin, Jeremy M. Bono, Helen K. Pigage, Carson W. Allan, Fernando Diaz, John R. McCoy, Clinton C. Green, Jeffrey B. Callan, Stephen P. Delahunt

## Abstract

Although RNA is found in the seminal fluid of diverse organisms, it is unknown whether this RNA is functional within females. Here, we develop an experimental proteomic method called VESPA (<u>V</u>ariant <u>E</u>nabled <u>S</u>ILAC <u>P</u>roteomic <u>A</u>nalysis) to test the hypothesis that *Drosophila* male seminal fluid RNA is translated by females. We find strong evidence for 67 male-derived, female-translated proteins (mdFTPs) in female lower reproductive tracts at six hours postmating, many with predicted functions relevant to reproduction. Gene knockout experiments indicate that genes coding for mdFTPs play diverse roles in postmating interactions, with effects on fertilization efficiency, and the formation and persistence of the insemination reaction mass, a trait hypothesized to be involved in sexual conflict. These findings advance our understanding of reproduction by revealing a novel mechanism of postmating molecular interactions between the sexes that strengthens and extends male influences on reproductive outcomes in previously unrecognized ways. Given the diverse species known to carry RNA in seminal fluid, this discovery has broad significance for understanding molecular mechanisms of cooperation and conflict during reproduction.

## Introduction

Reproductive success depends on complex molecular interactions between males and females that integrate morphological, physiological, and behavioral responses to mating^1,2^. Although fundamental to all sexually reproducing species, the mechanistic bases of many important postmating molecular interactions are still poorly understood^1,2^. Recently, it has become clear that males contribute more than just sperm to this process. Male seminal fluid is a complex mixture of diverse components, which not only aid in sperm survival and delivery to the oocyte but also interact directly with female reproductive tissues to facilitate fertilization and trigger lasting changes in female physiology and behavior^1–4^. While most attention has focused on the important roles played by seminal fluid proteins (SFPs) in driving the female postmating response, the functional significance of other seminal fluid components has received less attention^1,2,4–6^.

The presence of coding and noncoding RNA in sperm and seminal fluid is a common feature of male ejaculates, having been found in diverse organisms including mammals and insects^5,7–12^. While sperm RNA has received increasing attention and effects on developing offspring have been established, the functional significance of seminal fluid RNA remains unknown^5,13^. In some species, including humans, seminal fluid RNA is carried in extracellular vesicles (EVs) ^5,14^. The potential for EVs to serve as a mechanism for interorganismal signaling leads to the hypothesis that male RNA could be delivered to recipient cells in the female reproductive tract where it might perform critical functions^5,14^. The protein-coding potential of mRNA and diverse regulatory roles non-coding RNA plays in many fundamental biological processes^15^ suggest a novel mechanism by which males may mediate reproductive outcomes. This would have broad implications for understanding cooperative interactions between males and females that facilitate fertilization and antagonistic interactions within and between the sexes resulting from sexual selection and sexual conflict.

## Results

### Male seminal fluid RNA is translated by females

Building on our previous research showing the transfer of *Drosophila arizonae* male RNA to females during copulation^9^, here we test the hypothesis that male seminal fluid mRNA is translated into protein by females. Identification of male-derived, female-translated proteins (mdFTPs) requires not only differentiating male and female proteins within the female reproductive tract but also the source of transcripts from which these proteins were produced. To overcome this challenge, we developed VESPA (<u>V</u>ariant <u>E</u>nabled <u>S</u>ILAC <u>P</u>roteomic <u>A</u>nalysis), an experimental approach and bioinformatic pipeline that combines interspecies hybridization with metabolic proteome labeling of whole flies^16^ to identify mdFTPs (Fig. 1).

**Figure 1.**
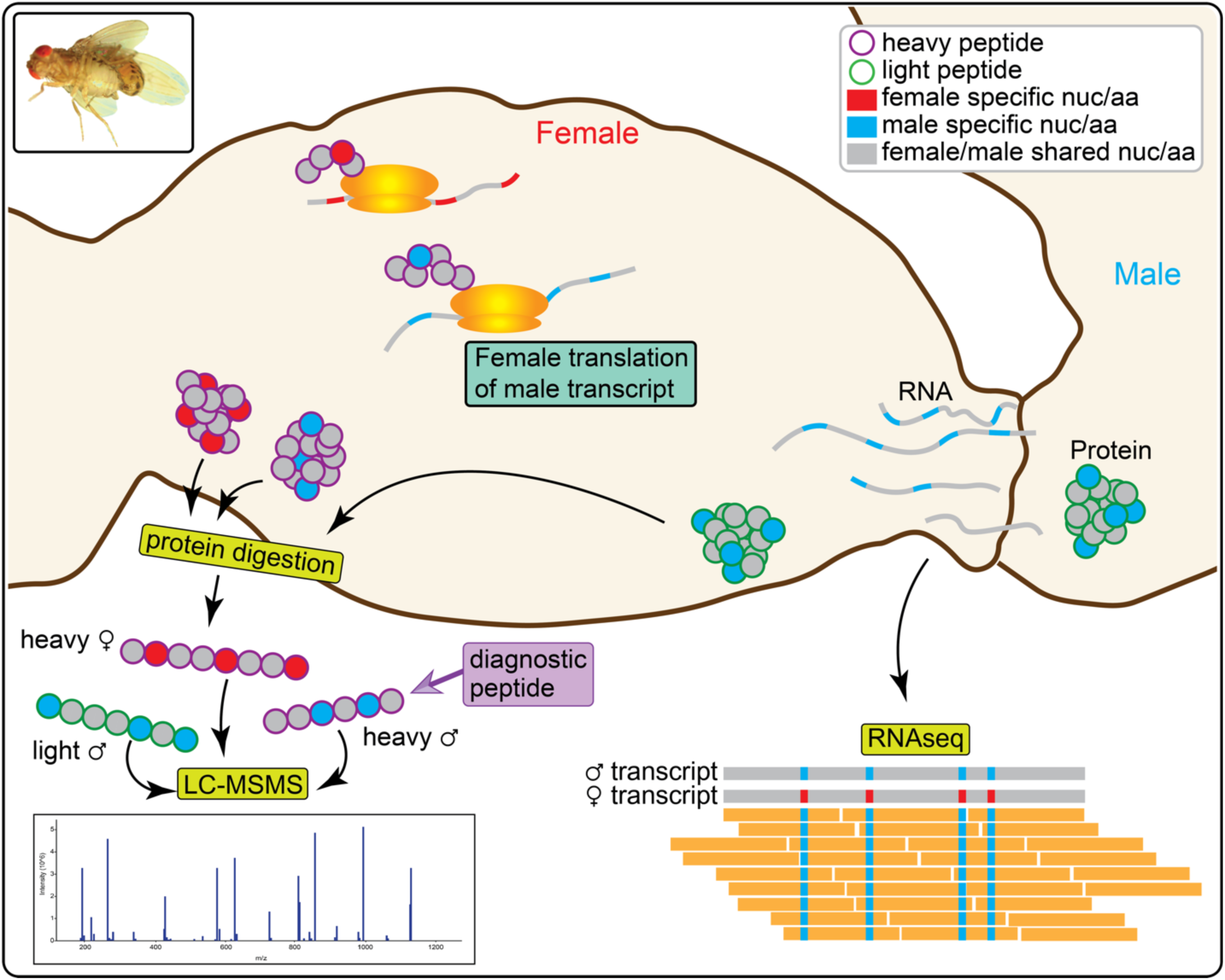
Overview of VESPA. VESPA differentiates proteins made by males and females by isotopic labeling, and the source of RNA transcripts by fixed nucleotide differences between species. Peptides that match the sequence of the male but carry the heavy label of the female are diagnostic for mdFTPs.

To identify mdFTPs using VESPA, we metabolically labeled the proteome of female *D. mojavensis* using L-Lysine-^13^C_6,_^15^N_2_ (Lys 8). Heavy labeled *D. mojavensis* females were mated with unlabeled *D. arizonae* males, which allows proteins produced by males and females to be differentiated within the female reproductive tract. While most peptide sequences are shared between the species, unique peptides resulting from fixed nonsynonymous substitutions distinguish the source of translated mRNA transcripts. With this approach, peptides that match the sequence of the male but the heavy isotope label of the female, are diagnostic for mdFTPs.

Lower reproductive tracts of heavy labeled *D. mojavensis* females mated to unlabeled *D. arizonae* males were removed at six hours postmating for proteomic analysis by Liquid Chromatography Tandem Mass Spectrometry (LC-MS/MS). The six hour timepoint was chosen based on previous results showing male RNA was still detectable in the female reproductive tract at this time (1), and because this would allow capture of proteins translated over the course of several hours. This experiment was independently repeated three times. Mass spectra were analyzed using MaxQuant^17^ and MSFragger^18^ with a combined database containing all annotated proteins in *D. mojavensis* and *D. arizonae*. These analyses resulted in a total of 290,733 (MaxQuant) and 889,134 (MSFragger) peptide spectrum matches (PSMs) across all three replicates using a false-discovery rate (FDR) threshold of 0.01. To find evidence for mdFTPs, we identified heavy-labeled PSMs that matched the *D. arizonae* database sequence (heavy-*arizonae*; HA PSMs). To further assess confidence in HA PSMs we compared features of these identifications with light-*mojavensis* (LM) PSMs. Since female proteomes were heavy labeled, LM PSMs were either erroneously identified or resulted from incomplete label incorporation. We identified an average of 7.2X more HA PSMs relative to LM PSMs across replicates (range: 4.1 – 9.1; TableS1), which further bolsters confidence in the overall validity of the set of HA PSMs. Output data from MaxQuant and MSFragger for diagnostic HA PSMs is included in Supplementary file 1.

After subsequent filtering to remove potential false positives resulting from polymorphism or leucine/isoleucine substitutions, a total of 234 and 187 unique HA peptides were identified by MaxQuant and MSFragger respectively (Table S2). To be considered a mdFTP, we required proteins to be identified by at least two heavy peptides (i.e. we filtered out so called ‘one-hit wonders’), one of which had to be a diagnostic HA peptide while the second could be diagnostic or non-diagnostic (peptide sequence shared between the species). This resulted in a total of 166 unique mdFTPs. Twenty-six mdFTPs were identified by multiple distinct HA peptides (range: 2-7), and 58 were identified in two or three replicates (Supplementary file 2). It is important to note that shotgun proteomics relies on a data-dependent acquisition strategy, where only a subset of peptides is selected for analysis. The stochastic nature of the selection process often results in peptides that are identified in only a subset of replicates^19,20^. Given that mdFTPs can only be identified by the relatively small number of diagnostic peptides that differ in sequence between *D. mojavensis* and *D. arizonae*, those with multiple diagnostic HA peptides, identified in multiple replicates, or by both search algorithms, have the strongest level of support (67 mdFTPs; Fig. 2A). Confidence in these mdFTPs is further strengthened by the high quality of associated diagnostic peptide identifications. On average, mdFTP diagnostic peptides had lower posterior error probabilities (PEP; MaxQuant) and higher peptide prophet confidence scores (MSFragger) relative to LM peptides, and these metrics were also similar to all other non-LM peptides (Fig. S1). The remaining mdFTPs were only identified in a single replicate, and, on average, diagnostic peptide confidence scores (PEP and peptide prophet probability) were similar to LM peptides and lower than highly supported mdFTP and other non-LM peptides (Fig S1). Nevertheless, these mdFTPs are supported by a median of seven heavy peptides total when including non-diagnostic peptides (Table S3). We conclude that while some mdFTPs in this group are likely valid identifications, the group as a whole may be enriched for false positives.

**Figure 2.**
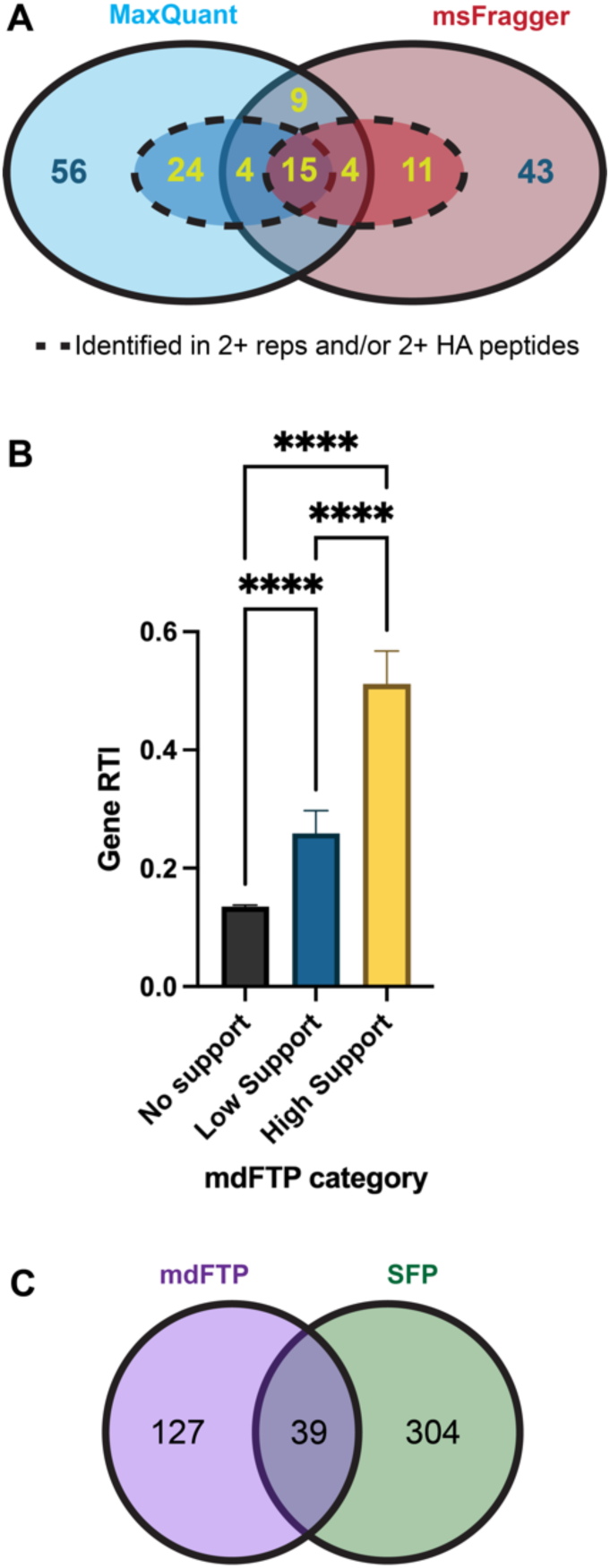
mdFTPs are supported by multiple lines of evidence. (A) All mdFTPs were identified by a minimum of two heavy peptides, with at least one being diagnostic. mdFTPs inside broken lines were identified in multiple replicates or by multiple diagnostic peptides. The most strongly supported mdFTPs include those identified by multiple diagnostic peptides, in two or more replicates, and/or by both MaxQuant and MSFragger. (B) Mean RNA transfer index (RTI) per gene was higher for high support mdFTPs (yellow in panel ‘A’, n=67) and low support mdFTPs (n=99) compared to all other genes with RNA-seq data (n=9727). (GLMM: gene category χ^2^=812.8, *P*= 2.2e^-16^). *Post hoc* tests were performed using Tukey’s method (\*\*\*\**P≤*0.0001). Error bars represent standard error of the mean. (C) Venn diagram showing overlap of mdFTPs with *D. arizonae* SFPs. Low overlap suggests mdFTPs may perform different functions from other proteins in the ejaculate.

### mdFTPs are supported by the presence of male RNA transcripts

mdFTPs are further supported by independent evidence from RNA-seq data. To investigate whether male RNA transcripts of mdFTPs were more likely to be detected in the reproductive tracts of heterospecifically mated females than transcripts of other proteins, we performed RNA-seq of *D. mojavensis* female lower reproductive tracts after mating with *D. arizonae* males. Although male transcripts can be identified by fixed nucleotide differences between the species, this is complicated by the fact that some male-transferred transcripts could be of low abundance and/or also be produced by females, resulting in observed variable sites in sequencing data. To evaluate evidence for male transcripts, we developed a metric called the RNA Transfer Index (RTI), which is the proportion of variable sites in a gene having at least five reads of male origin. Overall, transcripts of mdFTPs showed evidence of significant enrichment of male reads compared to non-mdFTPs. Specifically, mean RTI was ∼4X higher in the set of 67 highly supported mdFTPs, and ∼2X higher in the remaining set compared to non-mdFTPs (Fig. 2B). Transcripts of 11 mdFTPs appear to be supplied almost exclusively by males (>95% of reads are of male origin), while other mdFTP transcripts ranged from being mostly male in origin to mostly female. Overall, RNA-seq data provides strong support for a positive association between mdFTPs identified through proteomics and the presence of male RNA transcripts identified by RNA-seq in female lower reproductive tracts.

### mdFTPs are largely distinct from proteins transferred in the male seminal fluid

Male seminal fluid proteins (SFPs) play critical functional roles in the postmating response of diverse organisms^2,4^. To determine whether mdFTPs overlap with SFPs, we analyzed the proteomes of unlabeled *D. arizonae* female reproductive tracts immediately after copulation with Lys 4 labeled *D. arizonae* males. This analysis identified a total of 343 SFPs that are transferred to females during mating (Supplementary file 3). Only ∼25% overlapped with the overall set of mdFTPs (Fig. 2C), suggesting that most mdFTPs target distinct aspects of the postmating process or function through alternative pathways relative to SFPs.

### Functional clusters implicate mdFTPs in the postmating response

Gene ontology (GO) and protein domain enrichment analyses of the overall set mdFTPs revealed several enriched functional clusters with links to the postmating response (Fig. 3A). For example, proteins with a CAP domain were highly enriched. This domain is associated with cysteine-rich secretory proteins (CRISPs), which have diverse roles in fertility in mammals and insects^21–23^. GO-term analysis performed on mdFTP orthologs in *D. melanogaster* (126/166 mdFTPs had orthologous calls) revealed enrichment for processes related to metabolism and oxidative stress response. This is consistent with the fact that sperm must remain viable within the female reproductive tract potentially for days to weeks prior to fertilization. Additionally, one enriched functional cluster was related to immunity. The regulation of immunity by mating is a key component of the postmating response in both vertebrates and invertebrates^24,25^. Our data suggest a potentially novel mechanism by which female immunity is regulated by the male ejaculate. More detailed data on significant functional clusters is found in Supplementary File 4.

**Figure 3.**
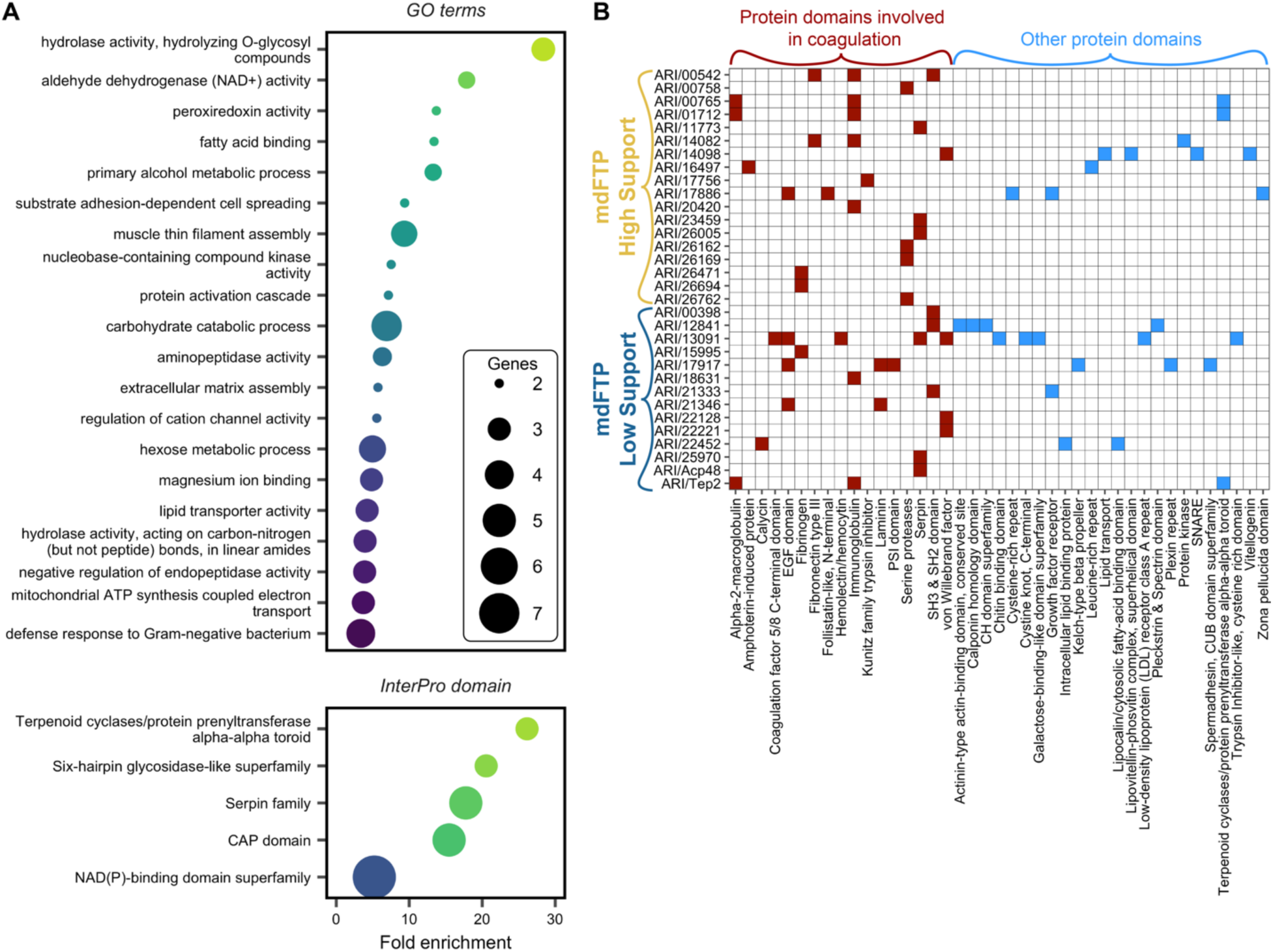
mdFTPs have functional significance to reproduction. (A) Enrichment of protein domains and GO terms for biological process and molecular function link mdFTPs to processes important for reproduction. This includes enrichment of CAP domains, which are linked to reproduction in diverse organisms, and two domains (serpin family and terpenoid cyclases/protein prenyltransferase alpha-alpha toroid) that have predicted involvement in coagulation. GO terms associated with energy production, oxidative stress response, and immunity were also enriched. (B) List of mdFTPs with protein domains associated with coagulation. Additional domains not necessarily linked to coagulation are also noted. High support mdFTPs refer to the 67 in yellow from figure 1A, while low support mdFTPs represent the remaining 99.

In *D. mojavensis* and *D. arizonae,* a large opaque mass called the insemination reaction mass forms in the female reproductive tract after mating^26,27^. The composition of the reaction mass is unknown, but it is hypothesized to mediate sexual conflict over female remating rate^26,28^. In conspecific crosses, it persists for several hours, but in crosses between *D. mojavensis* and *D. arizonae* it lasts much longer, potentially even permanently sterilizing females^26,28^. Notably, at least two mdFTPs, ARI/13091 (CactusFlybase id: CFgn0008961; https://cactusflybase.arizona.edu/) and ARI/14098 (CFgn0013989) orthologous to *D. melanogaster hml* and *apolpp* respectively, have demonstrated roles in hemolymph clotting in *Drosophila*^29,30^. This finding is especially interesting given that the reaction mass physically resembles a coagulatory response. In addition, two enriched protein domains, serpin family and terpenoid cyclases/protein prenyltransferase alpha-alpha toroid, which overlaps with alpha-2 macroglobulin, have also been implicated in clotting and wound healing in vertebrates and some invertebrates^31–33^. Overall, at least 32 mdFTPs contain conserved protein domains associated with clotting and wound healing (Fig. 3B), particularly in vertebrates^34^.

### mdFTPs are involved in formation and degradation of the reaction mass

Based on the large number of mdFTPs associated with coagulation, we hypothesized that the reaction mass is caused by a coagulatory response induced by SFPs and/or mdFTPs. Among the proteins linked to clotting, three mdFTPs contain fibrinogen domains, and males transfer several other fibrinogens as SFPs (Fig. 3B; Supplementary File 3). In vertebrates, fibrinogens are a critical component of the blood clotting cascade^35^, though invertebrate fibrinogens are predicted to have other functions not associated with coagulation^36^. To test the hypothesis that fibrinogens are involved in the formation of the reaction mass, we used CRISPR to knock out the gene *ARI/26694* (CFgn0007404), which is a fibrinogen from the highly supported set of mdFTPs that is transferred as both mRNA and protein to females. We compared the size of the reaction mass in females mated to KO or wild-type (WT) males immediately after mating and at six hours postmating. Females mated to KO males initially had a smaller reaction mass and the mass also degraded more slowly compared to females mated to WT males (Fig. 4A; Fig. S2). Although we currently cannot differentiate the contribution of protein transferred by the male from protein produced from male RNA by the female, this result indicates that genes coding for mdFTPs are involved in a key aspect of the postmating response with predicted implications for sexual conflict.

**Figure 4.**
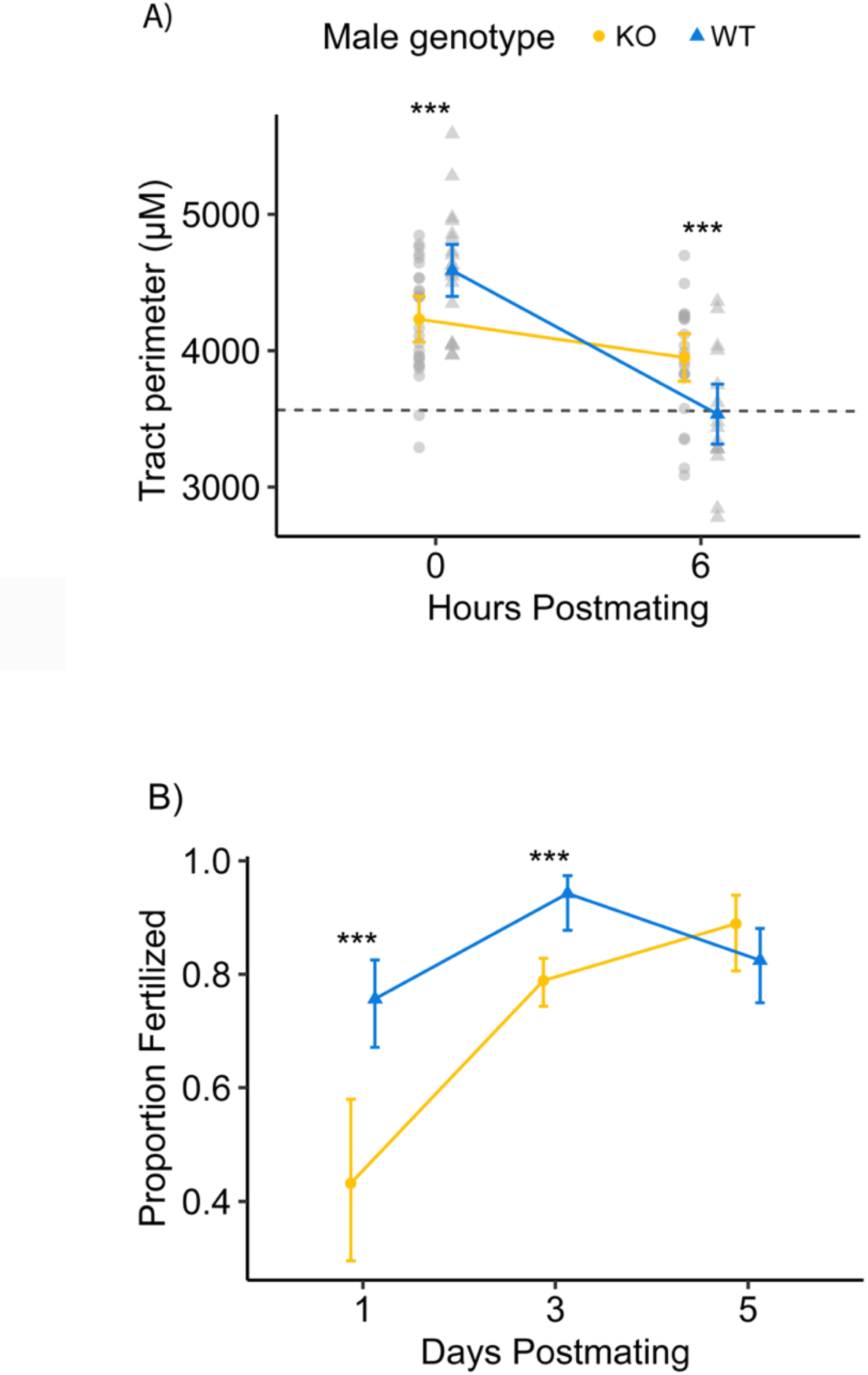
Gene KO experiments demonstrate mdFTPs have functional effects on diverse reproductive processes. (A) *Drosophila arizonae* females mated to *D. arizonae* males with a *ARI26694* KO mutation had a smaller reaction mass initially, as measured by the average perimeter. However, it degraded more slowly compared to the reaction mass in females mated to WT males (Two-way ANOVA: genotype x time interaction, F= 16.5, *P*= 0.0001). *Post hoc* comparisons were performed using Tukey’s method (\*\**P≤*0.01). The dashed line indicates the mean perimeter of unmated female lower reproductive tract. Error bars represent 95% confidence intervals of estimated marginal means. KO-0 hr: n=26; WT-0 hr: n=35; KO-6 hr: n=15. (b) *D. arizonae* females mated to *D. arizonae* males with a *ARI11629* KO mutation laid more unfertilized eggs on days one and three postmating compared to females mated to WT males (GLM: genotype x day interaction, χ^2^= 17.2, *P*= 0.0002). *Post hoc* comparisons were performed using Tukey’s method (\*\*\**P*> 0.001). Error bars represent 95% confidence intervals of estimated marginal means. WT-D1: n=119; KO-D1: n=44; WT-D3: n=104; KO-D3 n=360; WT-D5 n=131; KO-D5 n=90.

### mdFTPs affect fertilization efficiency

To evaluate whether mdFTPs have broader effects on postmating responses beyond the reaction mass, we used CRISPR to knockout the gene *ARI/11629* (CFgn0010028). This cysteine peptidase is one of the most strongly supported mdFTPs, being identified by four diagnostic peptides in multiple replicates across both search programs, and with evidence for male RNA transcripts in this and a previous study^9^. This gene is also evolving rapidly in *D. mojavensis* and *D. arizonae*^37^, which is consistent with involvement in sexual selection and/or sexual conflict. We compared eggs laid over seven days and fertilization efficiency for WT females mated to WT or KO males. While females mated to KO and WT males laid a similar number of eggs (Fig. S3), fertilization efficiency in females mated to KO males was markedly reduced at days one and three postmating (Fig. 4B). Since this protein is also transferred in the seminal fluid, additional studies are necessary to tease apart the effect protein supplied by the male from protein translated by females from male RNA. Nevertheless, these results indicate genes coding for mdFTPs have broad-ranging effects on the postmating response that extend for days after mating.

## Discussion

The results of this study advance our understanding of the molecular mechanisms of reproduction by illuminating a previously unrecognized mode of postmating interaction between males and females. One of the main benefits of transferring RNA to females may be that it provides a mechanism to temporally extend the production of key proteins, as most male proteins are degraded within a few hours in the highly proteolytic environment of the female reproductive tract^38–40^. Transcriptome data from this study and our previous study^9^ indicate male RNA persists within the female reproductive tract for at least six hours postmating (longer time points have not been tested). Temporal control over protein production could be beneficial given that sperm must remain viable and compete with sperm from other males for days to weeks after mating. Recent evidence in *D. melanogaster* demonstrates that a “molecular handoff” between the sexes occurs within the female reproductive tract, whereby females gradually take over the production of key sperm proteins initially supplied by males^40^. These proteins are enriched for processes such as energy metabolism, which is assumed to aid in sperm survival^40^. This type of cooperative maintenance of sperm viability between the sexes is predicted to be widespread across animals^40^, but the molecular mechanisms of how the molecular handoff occurs have not been elucidated. Orthologs of at least 12 mdFTPs overlap with this set of female-derived sperm-associated proteins in *D. melanogaster* ^40^. Moreover, mdFTPs are highly enriched for processes associated with energy metabolism and oxidative stress response. This suggests that molecular continuity between the sexes may be facilitated, at least in part, by the intersexual RNA transfer described here.

Another benefit of transferring RNA to females is that transcripts could be targeted to specific cells in the female reproductive tract where they could be translated. Localization of protein within female cells could provide functional capabilities that would not be possible for proteins transferred to the lumen of the female reproductive tract, and repeated translation from RNA could generate higher quantities of protein. Cell targeting could be facilitated if RNA is carried in extracellular vesicles (EVs), which are capable of targeting cargo to specific cell types. We do not know how *D. arizonae* male seminal fluid RNA is packaged, but previous studies in *D. melanogaster* have shown that males transfer exosomes to females during copulation^41^. Although the contents of these exosomes have not been identified, they fuse with female epithelial cells and sperm and have demonstrated effects on female postmating physiology and behavior^41–43^. These findings, coupled with the fact that seminal fluid RNA in other organisms is packaged in EVs^5,14^, suggests this possibility warrants further study.

Formation of an insemination reaction mass is relatively rare across *Drosophila* species^26^, and the molecular bases of the formation and degradation of the mass are unknown. We found that ∼30% of mdFTPs overlapped with the set of over 1500 proteins associated with the *D. melanogaster* mating plug^44^, which is structurally distinct from the reaction mass. mdFTPs also include two proteins with known roles in *D. melanogaster* hemolymph coagulation (*hml* and *apolpp*) ^29,30^, which is consistent with the clot-like appearance of the reaction mass. However, other proteins known to play crucial roles in hemolymph coagulation and mating plug formation were not observed among mdFTPs or SFPs^29,30^. These findings suggest that the reaction mass is formed through distinct molecular pathways relative to other coagulatory responses in *Drosophila.* Interestingly, mdFTPs include representatives of several protein classes associated with blood clotting in vertebrates^34^, but that have not been linked to clotting in invertebrates^29,30^ (Fig. 3B). Some of these protein classes have broad functional roles, including other functions related to reproduction. Nevertheless, the confluence of all of them, along with the presence of the reaction mass, is striking. Our results demonstrate that *ARI/26694*, a fibrinogen, is involved in the formation and degradation of the reaction mass. While vertebrate fibrinogens play a critical role in blood clotting, previous studies have suggested that invertebrate fibrinogens do not function in coagulation^36^. Altogether, our findings suggest the reaction mass forms through a coagulatory response that is influenced in part by genes coding for mdFTPs.

The reaction mass is hypothesized to serve as a mating plug that delays female remating ^26,28^, although other functions are also possible^45,46^. While recent studies in insects have revealed female contributions to the formation or degradation of mating plugs^44,47^, our results suggest males induce females to participate in the process of generating and/or degrading the reaction mass through the transfer of RNA. While it may seem contradictory that *ARI/26694* is involved in both the formation and degradation of the reaction mass, vertebrate fibrinogens are known to facilitate the assembly and degradation of fibrin clots^48^. Consistent with this, the set of mdFTPs with predicted roles in clotting include proteins with pro- and anticoagulatory activity (Fig. 3B). Notably, while the reaction mass may prevent rapid remating, it also delays the onset of oviposition^28,49^. Thus, males may benefit from playing a role in both the formation and dissolution of the mass. A similar mechanism is observed in primates, where males transfer proteins to females that are involved in both the formation and degradation of a copulatory plug^50^. Overall, these findings indicate that the formation and degradation of the reaction mass is a highly orchestrated process that might be facilitated by the transfer of male RNA to females during copulation. If future studies confirm a direct role for mdFTPs in this process, this could have broad implications for understanding the formation and degradation of mating plugs, which have evolved independently in many taxa.

## Conclusions

Sexual reproduction is a complex process that involves both cooperation and conflict between the sexes at the molecular level. Here, we illuminate a previously unrecognized mode of molecular interaction and interdependence between males and females. We show that males functionally alter the female postmating proteome by transferring RNA during copulation that is subsequently translated by females. Moreover, we demonstrate that genes coding for mdFTPs have diverse influences on postmating outcomes including effects on fertilization efficiency and formation/degradation of the reaction mass, a trait with predicted involvement in sexual conflict. Given the powerful coding and regulatory properties of RNA, coupled with the fact that RNA is a common feature of male ejaculates, this discovery has important implications for understanding the molecular mechanisms of reproduction across the tree of life.

## Material and methods

### Drosophila stocks

All proteomic experiments described used *D. mojavensis* females (National *Drosophila* Species Stock Center (NDSSC): 15081-1353.01) and *D. arizonae* males (NDSSC: 15081-1271.41). Genomes from these stocks are accessible at the cactusflybase site (https://cactusflybase.arizona.edu/). KO *D. arizonae* lines were also generated in the same *D. arizonae* stock.

### Metabolic labeling of Drosophila

To distinguish the origin of proteins in the female reproductive tract, we metabolically labeled female flies, by modifying protocols developed previously for *D. melanogaster* ^16,40,51–54^. Males were unlabeled. For labeling, we reared a lysine auxotrophic strain of *Saccharomyces cerevisiae* (MATalpha leu2Δ0 lys2Δ0 ura3Δ0 in background BY4729; Dharmacon, Inc. Lafayette, CO) in synthetic media containing yeast nitrogen base without amino acids, yeast synthetic drop-out medium without lysine, and isotopically labeled lysine (L-Lysine-^13^C_6,_^15^N_2_ or Lys8; Cambridge Isotopes Laboratory, Inc, Tewksbury, MA). Cultures were grown in a shaking incubator for ∼24 hours at 30°C. Cells were pelleted by centrifugation, washed with sterilized water, lyophilized, and frozen at -20°C. We made food media for rearing flies by combining Lys8 labeled yeast, yeast nitrogen base without amino acids, pure molasses, agar, sterile water and methylparaben dissolved in ethanol as a preservative (Table S4). This mixture was autoclaved and dispensed into sterile glass vials in a biosafety cabinet. We placed *D. mojavensis* adults in population cages overnight on media containing baker’s yeast and molasses to encourage oviposition. The following day, eggs were collected and sterilized/dechorionated by soaking in 2.5% sodium hypochlorite for three minutes. Eggs were rinsed with sterile water and transferred under aseptic conditions to vials containing Lys8 food (∼100 eggs/vial) where they were reared until adult emergence. We collected unmated female flies within a few hours of eclosion and transferred them to vials containing sterilized food consisting of pure molasses, agar, water, and a flake of lyophilized Lys8 labeled yeast. Flies were transferred daily to new vials until they reached reproductive maturity. This protocol ensured that flies were only exposed to Lys8 labeled protein prior to mating experiments, resulting in high labeling efficiency (∼99%). Heavy-labeled unmated female *D. mojavensis* were paired with unmated *D. arizonae* males that had been reared in standard banana-molasses food in vials containing sterilized molasses-agar food without yeast^55^. Copulations were observed and mated females were isolated until we removed lower reproductive tracts at six hours postmating. Female reproductive tracts were placed in 20 μl ice cold 50mM ammonium bicarbonate. Individual tubes were moved to -80°C once five tracts were collected. This experiment was repeated three times, with each biological replicate consisting of ∼45 lower reproductive tract.

### Protein isolation and LC-MS/MS

Protein isolation and LC-MS/MS was performed at the Central Analytical Mass Spectrometry Facility and W.M. Keck Foundation Proteomics Resource at University of Colorado Boulder. Heterospecifically-mated *D. mojavensis* lower female reproductive tracts were denatured, reduced and alkylated with 5% (w/v) sodium dodecyl sulfate (SDS), 10mM tris (2-carboxyethyl) phosphine hydrochloride (TCEP-HCl), 40mM 2-chloroacetamide, 50mM Tris pH 8.5 and boiled at 95°C for 10 minutes. Samples were prepared for mass spectrometry analyses using the SP3 method^56^. Carboxylate-functionalized speedbeads (Cytiva) were added to protein lysates. Acetonitrile was added to 80% (v/v) to precipitate protein and bind it to the beads. The protein-bound beads were washed twice with 80% (v/v) ethanol and twice with 100% acetonitrile. LysC/Trypsin mix (Promega) was added for approximately 1:50 protease to protein ratio in 50mM Tris pH 8.5 and incubated rotating at 37°C overnight. Tryptic digests were cleaned using an Oasis HLB 1cc (10mg) cartridge (Waters) according to the manufacturer. Samples were dried using a vacuum rotatory evaporator. To reduce sample complexity, samples were fractionated using a Waters M-class UPLC equipped with a photodiode-array detector. Samples were suspended in 50uL 0.16% (v/v) aqueous ammonia in water, then injected onto a 500um id x 150mm long custom fabricated rpC18 column (UChrom C18 1.8um 120A; www.nanolcms.com) and separated with a gradient 2% to 40% acetonitrile with 0.16% (v/v) aqueous ammonia. Concatenated fractions (8-12 depending on the replicate) were collected and dried using a vacuum rotatory evaporator. Fractions were suspended in 0.1% TFA, 3% acetonitrile in water for LC/MS/MS analyses. Fractionated peptides were directly injected onto a Waters M-class column (1.7um, 120A, rpC18, 75um x 250mm) and gradient eluted from 2% to 20% acetonitrile with 0.1%(v/v) formic acid over 100 minutes then 20% to 32% acetonitrile over 20 minutes at 0.3uL/minute using a Thermo Ultimate 3000 UPLC (Thermo Scientific). Peptides were detected with a Thermo Q-Exactive HF-X mass spectrometer (Thermo Scientific) scanning MS1 at 120,000 resolution from 380 to 1580 m/z with a 45ms fill time and 3E6 AGC target. The top 12 most intense peaks were isolated with 1.4 m/z window with a 100ms fill time and 1E5 AGC target and 27% HCD collision energy for MS2 spectra scanned at 15,000 resolution. Dynamic exclusion was enabled for 25 seconds.

### Identification of mdFTPs

Mass spectra were analyzed using two different software programs: MaxQuant^17^ (ver. 2.0.3.0) and MSFragger^18^ (ver. 3.6 within FragPipe ver. 19.0). Database searches assumed SILAC Lys8 labeling, and also included common variable modifications (Oxidation (M); Acetyl (Protein N-term)). PSM and protein FDR were set to 0.01. The ‘requantify’ and ‘matching between runs’ features were not utilized. The protein database for both *D. mojavensis* and *D. arizonae* was based on our recent assembly and annotation including all genome transcripts (19,778 for *D. mojavensis* and 19,747 for *D. arizonae*)^57^ available at OSF. All gene IDs used in this study follow annotation nomenclature set forth in the cactusflybase site^57^. PSMs were sorted to identify heavy *D. arizonae* (HA) PSMs, which are diagnostic for mdFTPs. We required mdFTPs to be supported by a minimum of one diagnostic peptide and at least one additional heavy peptide that could be diagnostic or non-diagnostic (a heavy peptide with the same sequence in *D. arizonae* and *D. mojavensis*). Thus, so called ‘one-hit-wonders’ were filtered out of our mdFTP list. All mass spectrometry proteomics data have been deposited on the ProteomeXchange Consortium via the PRIDE^58^ partner repository with the dataset identifier PXD041195. All MaxQuant and MSFragger analysis parameter and output files are available at OSF.

### HA peptide filtering and quality assessment

We performed rigorous filtering to remove potential false positive HA peptide identifications due to polymorphism or leucine/isoleucine substitutions. Moreover, to further investigate the validity of HA peptide identifications we compared features of HA peptides with LM peptides, which could only be identified by error or because of incomplete label incorporation.

The consensus genome sequences of *D. mojavensis* and *D. arizonae* do not include potential polymorphisms that could confound identification of diagnostic peptides (e.g. one *D. mojavensis* segregating allele matches *D. arizonae*). Therefore, we collected the head and thorax of mated females from one of our experimental replicates and bulk sequenced the sample so that any HA peptides identified based on polymorphic sites could be filtered out. DNA from the head and thorax of 45 mated females was extracted using Qiagen DNeasy Blood & Tissue Kit (Qiagen, Hilden, Germany). The library was prepared using the KAPA LTP Library Preparation Kit (Roche, Basel, Switzerland) kit and sequenced on an Illumina HiSeq 4000 at Novogene (Beijing, China) to 114X coverage. Raw reads have been deposited on NCBI’s SRA repository under BioProject PRJNA949702. Reads from *D. mojavensis* were trimmed using Trimmomatic^59^ and mapped to the Sonora *D. arizonae* genome r0.93^57^ using bwa-mem^60^. PCR duplicates were then removed using Picard v2.27.5^61^.

To determine if polymorphism existed within diagnostic peptides, we first mapped HA peptides to the *D. arizonae* genome r0.93^57^ using ACTG^62^. To utilize ACTG, we generated a GFF file from the GTF using AGAT^63^. From the output, a BED file was generated of all HA peptides, and we used bedtools2^64^ to extract the peptide regions. A mpileup file was then generated using samtools^65^, and variant calling was performed with VarScan2^66^. SnpEff^67^ was used to identify synonymous and nonsynonymous changes in the candidate peptide regions. We retained peptides if they met the following two conditions: (1) at least one nonsynonymous change that was not a Leu-Ile or Ile-Leu (not distinguishable by mass spectrometry), (2) bulk genome sequencing of *D. mojavensis* revealed that the sequence was not polymorphic in *D. mojavensis* (≥95% of reads match the *D. mojavensis* reference genome).

To further investigate the validity of HA peptide identifications we compared the number of HA PSMs to LM (light-*mojavensis*) PSMs identified in each replicate. Since female proteomes were heavy labeled, any LM PSMs are assumed to be erroneous identifications (we excluded PSMs including both heavy and light quantifications since they could represent proteins with incomplete label incorporation). HA PSMs could be errors or could represent mdFTPs. Since LM and HA PSMs should be equally likely to be erroneously identified, we expect the number of each type to be approximately equal if most HA PSMs are errors.

### RNA-seq of mated female reproductive tracts

We used RNA-seq to identify the presence of male-derived transcripts in the reproductive tracts of *D. mojavensis* females mated to *D. arizonae* males. We used a combination of DNA and RNA sequencing to establish a set of fixed nucleotide differences between *D. mojavensis* and *D. arizonae* since consensus genome sequences do not provide information on polymorphism within lines. For *D. arizonae*, we identified polymorphisms using genome sequencing reads and pooled RNA-seq reads from multiple life stages originally collected for the *D. arizonae* genome assembly^57^. For *D. mojavensis*, we used reads from the genome assembly and from head and thorax samples of mated flies described above. Reads from both species were mapped to the *D. arizonae* r0.93 assembly^57^ using HISAT2^68^ for RNA and bwa^60^ for DNA. Duplicates were then removed using Picard v2.27.5^61^, bam files were merged, exons regions exported, mpileup generated using samtools^65^ and variants called using VarScan2^66^. Variant called files for *D. arizonae* DNA and RNA sequencing were generated containing positions which had a *D. mojavensis* allele frequency > 0.95 and a significant Fisher’s Exact test (P < 0.05). We used these files to generate a list of 549,579 positions that represent fixed differences between the species.

To identify male derived RNA in the lower reproductive tract of females, we used RNA-seq data from^69^. Reads from heterospecifically-mated *D. mojavensis* females 45 minutes and 6 hours postmating (NCBI BioProject PRJNA777940) were trimmed with Trimmomatic^59^, duplicates removed with Picard v2.27.5^61^, and pooled and mapped to the *D. arizonae* r0.93 genome using HISAT2^68^. We used the list of fixed sites between the species to query the RNA-seq data to detect male reads (*D. arizonae*). Using a custom perl script, we determined the frequency of the male allele at each site. This was then used to calculate the RNA transfer index (RTI), which is the proportion of sites per gene that have 5 or more *D. arizonae* reads. We compared the RTI among three categories of genes: highly supported mdFTPs, supported mdFTPs, and genes with no support for being mdFTPs. Comparisons were made using a generalized linear mixed model (GLMM) with binomial error distribution implemented in the ‘lme4’^70^ package for R. Gene category was treated as a fixed effect and gene id was treated as a random effect. An anova table was generated using the ‘car’^71^ package, and *post-hoc* testing with Tukey’s adjustment was performed using the ‘multcomp’^72^ package.

### Identification of D. arizonae male seminal fluid proteins

*Drosophila arizonae* SFPs were identified as part of a larger unpublished analysis comparing *D. mojavensis/D.arizonae* seminal fluid proteomes. *Drosophila arizonae* were reared in standard banana-molasses food^55^ or food made with L-lysine-2HCL,4,4,5,5-D4 (Lys 4; Cambridge Isotopes Laboratory, Inc, Tewksbury, MA) as described above. Unmated Lys 4 males and unlabeled females were paired in vials and copulations were observed. Mated females were immediately frozen in liquid nitrogen and stored at -80°C until reproductive tracts were removed and placed in 50 mM ammonium bicarbonate. These tracts were combined in groups of five with mated *D. mojavensis* tracts that were collected in the same manner, except males were labeled with Lys 8 (six biological replicates). Protein extraction, digestion, LC-MS/MS, and analysis with MaxQuant^17^ (ver. 2.0.3.0) were performed as described above. *Drosophila arizonae* male SFPs were identified by filtering the protein list to include only proteins carrying the Lys 4 label. We considered identified proteins to be SFPs if they were found in any of the six replicates.

### GO-term and protein domain enrichment analyses

To obtain functional predictions for mdFTPs we analyzed GO-terms and protein domains. GO-term enrichment was analyzed using ClueGO^73^. Since GO-terms are better annotated for *D. melanogaster,* we used *D. melanogaster* orthologs of mdFTPs (126/167 had orthologous calls). The background gene list for gene enrichment analysis included orthologs of all *D. arizonae* genes identified in *D. melanogaster*. We tested for enrichment in terms for biological process and molecular function using a right-sided hypergeometric test applying a Benjamini-Hochberg false-discovery rate threshold of 0.05. Enriched terms were grouped by ClueGO based on functional relationships. We report the average fold enrichment per group. To assess protein domain enrichment, we first identified domains in the *D. arizonae* genome using InterProScan^74^ to generate a database. We used the R package WebGestalR^75^ to determine overrepresentation of domains in our mdFTPs using the *D. arizonae* as its background.

### Knockout experiments

We used CRISPR knockouts to test the effects of *ARI/26694* and *ARI11629* on aspects of the female postmating response. Genes were targeted using two sgRNAs designed against regions at the 5’ end of each gene following the method outlined by Bassett et al.^76^ sgRNAs and Cas9 mRNA (TriLink Biotechnologies, San Diego, CA) were injected into *D. arizonae* embryos by Rainbow Transgenic Flies, Inc. (Camarillo, CA). For each gene, we generated homozygous lines for two different frameshift mutations (Fig. S4), which were crossed to make transheterozygous males used in experiments. On the morning of experiments, unmated WT females from a different *D. arizonae* line (ARTU2) were paired with either unmated WT or KO males and copulations were observed. For *ARI/26694*, females were separated from males and placed in liquid nitrogen either immediately after mating or six hours postmating. Lower reproductive tracts were removed and photographed using a camera mounted to a Leica S9i dissecting microscope (37.5X magnification). Images were analyzed using ImageJ^77^ software by using the freeform drawing tool to trace the outline of the lower female reproductive tract. Perimeter and area were calculated, and pixels were converted to μm using photo resolution. Data were analyzed as a factorial type II Anova using the ‘afex’^78^ package for R. The model included male genotype, time, and their interaction as factors. Model fit was evaluated visually using residual plots. *Post-hoc* comparisons were analyzed using the ‘emmeans’^79^ package for R with Tukey’s adjustment for multiple comparisons.

For *ARI/11629*, we performed two experiments analyzing egg-laying and fertilization efficiency. For egg-laying, females were separated from males after copulation and placed alone in vials containing banana-molasses food. Flies were moved to new vials every 24 hours for seven days and eggs were counted each day. Data were analyzed with the R package ‘glmmTMB’^80^ using a generalized linear mixed model (GLMM) with a negative binomial error distribution. The model included male genotype, day, and their interaction. For the fertilization efficiency experiment, mated females were separated from males and placed in population cages overnight on banana-molasses food with yeast paste. The food plate was removed in the morning and stored for 6 h so that developing embryos would be 6 to 22 h. Embryonic development time for these species is approximately 28 hours^81^. We used a solution of 2.5% sodium hypochlorite to dechorionate embryos before they were fixed in a of 1:4 solution of 4% paraformaldehyde/heptane and then devitellinized in methanol. We stained embryos with 4′,6-diamidino-2-phenylindole (DAPI; 2.8 μg/ml) and analyzed embryonic development by fluorescence microscopy using a Leica DM5000B microscope (20X objective). We considered eggs to be unfertilized if no more than four nuclei were observed (representing the four products of female meiosis). We acknowledge that although the minimum age of embryos was six hours, embryogenesis could have been arrested at an earlier stage of development if the embryo was inviable. If this occurred during the earliest syncytial divisions, these embryos would be difficult to distinguish from unfertilized eggs using our methodology. However, this possibility is unlikely given that we did not observe embryos at a range of development times between the earliest syncytial divisions and 6 hours, as would be expected if inviability was common. Moreover, we would expect inviability to remain constant over time, but the phenotype we observed changed over time.

## Acknowledgements

We thank Chris Ebmeier from the Central Analytical Mass Spectrometry Facility and W.M. Keck Foundation Proteomics Resource at University of Colorado Boulder for assistance with proteomic analyses. We also thank Todd Schlenke for constructive comments on the manuscript.

## Data availability

Raw sequence data reported has been deposited in NCBI SRA under accession number PRJNA949702. Proteomic data has been deposited in PRIDE under project ID PXD041195. Remaining datasets and analysis files can be accessed via OSF. All fly lines generated for this study are available upon request from the corresponding authors. Code for analysis pipeline is included in supplementary information.

## References

1. Pitnick, S., Wolfner, M. F. & Dorus, S. Post-ejaculatory modifications to sperm (PEMS). Biological Reviews 95, 365–392 (2020).

2. Pitnick, S., Wolfner, M. F. & Suarez, S. S. Ejaculate-female and sperm-female interactions. in Sperm Biology (eds. Birkhead, T., Hosken, D. & Pitnick, S.) 247–304 (Elsevier Ltd, 2009). doi:10.1016/B978-0-12-372568-4.00007-0.

3. Poiani, A. Complexity of seminal fluid: A review. Behavioral Ecology and Sociobiology vol. 60 289–310 (2006).

4. Schjenken, J. E. & Robertson, S. A. The female response to seminal fluid. Physiol Rev 100, 1077–1117 (2020).

5. Jodar, M. Sperm and seminal plasma RNAs: What roles do they play beyond fertilization? Reproduction vol. 158 R113–R123.

6. Scolari, F., Khamis, F. M. & Pérez-Staples, D. Beyond Sperm and Male Accessory Gland Proteins: Exploring Insect Reproductive Metabolomes. Front Physiol 12, 1716 (2021).

7. Fischer, B. E. et al. Conserved properties of *Drosophila* and human spermatozoal mRNA repertoires. Proceedings of the Royal Society B: Biological Sciences 279, 2636–2644 (2012).

8. Lalancette, C., Miller, D., Li, Y. & Krawetz, S. A. Paternal contributions: New functional insights for spermatozoal RNA. Journal of Cellular Biochemistry vol. 104 1570–1579 (2008).

9. Bono, J. M., Matzkin, L. M., Kelleher, E. S. & Markow, T. A. Postmating transcriptional changes in reproductive tracts of con- and heterospecifically mated *Drosophila mojavensis* females. Proc Natl Acad Sci U S A 108, 7878–7883 (2011).

10. Degner, E. C. et al. Proteins, transcripts, and genetic architecture of seminal fluid and sperm in the mosquito *Aedes aegypti*. Molecular and Cellular Proteomics 18, S6–S22 (2019).

11. Ahmed-Braimah, Y. H., Wolfner, M. F. & Clark, A. G. Differences in Postmating Transcriptional Responses between Conspecific and Heterospecific Matings in *Drosophila*. Mol Biol Evol 38, 986–999 (2021).

12. Alfonso-Parra, C. et al. Mating-Induced Transcriptome Changes in the Reproductive Tract of Female *Aedes aegypti*. PLoS Negl Trop Dis 10, (2016).

13. Santiago, J., Silva, J. v., Howl, J., Santos, M. A. S. & Fardilha, M. All you need to know about sperm RNAs. Hum Reprod Update 28, 67–91 (2021).

14. Tamessar, C. T. et al. Roles of male reproductive tract extracellular vesicles in reproduction. American Journal of Reproductive Immunology 85, e13338 (2021).

15. Cech, T. R. & Steitz, J. A. The noncoding RNA revolution-trashing old rules to forge new ones. Cell 157, 77–94 (2014).

16. Sury, M. D., Chen, J. X. & Selbach, M. The SILAC fly allows for accurate protein quantification in vivo. Molecular and Cellular Proteomics 9, 2173– 2183 (2010).

17. Tyanova, S., Temu, T. & Cox, J. The MaxQuant computational platform for mass spectrometry-based shotgun proteomics. Nature Protocols 2016 11:12 11, 2301–2319 (2016).

18. Kong, A. T., Leprevost, F. v, Avtonomov, D. M., Mellacheruvu, D. & Nesvizhskii, A. I. msFragger: ultrafast and comprehensive peptide identification in mass spectrometry-based proteomics. 14, 513 (2017).

19. Liu, H., Sadygov, R. G. & Yates, J. R. A model for random sampling and estimation of relative protein abundance in shotgun proteomics. Anal Chem 76, 4193–4201 (2004).

20. Tabb, D. L. et al. Repeatability and Reproducibility in Proteomic Identifications by Liquid Chromatography—Tandem Mass Spectrometry. J Proteome Res 9, 761 (2010).

21. Gonzalez, S. N., Sulzyk, V., Weigel Muñoz, M. & Cuasnicu, P. S. Cysteine-Rich Secretory Proteins (CRISP) are Key Players in Mammalian Fertilization and Fertility. Front Cell Dev Biol 9, 3438 (2021).

22. Laflamme, B. A. & Wolfner, M. F. Identification and Function of Proteolysis Regulators in Seminal Fluid. Mol Reprod Dev 80, 80 (2013).

23. Avila, F. W., Sirot, L. K., Laflamme, B. A., Rubinstein, C. D. & Wolfner, M. F. Insect seminal fluid proteins: Identification and function. Annu Rev Entomol 56, 21–40 (2011).

24. Lawniczak, M. K. N. et al. Mating and immunity in invertebrates. Trends in Ecology and Evolution vol. 22 48–55 (2007).

25. Morrow, E. H. & Innocenti, P. Female postmating immune responses, immune system evolution and immunogenic males. Biological Reviews vol. 87 631–638 (2012).

26. Alonso-Pimentel, H., Tolbert, L. P. & Heed, W. B. Ultrastructural examination of the insemination reaction in *Drosophila*. Cell Tissue Res 275, 467–479 (1994).

27. Markow, T. A. & Ankney, P. F. Insemination Reaction in *Drosophila*: Found in Species Whose Males Contribute Material to Oocytes Before Fertilization. Evolution (N Y) 42, 1097 (1988).

28. Knowles, L. L. & Markow, T. A. Sexually antagonistic coevolution of a postmating-prezygotic reproductive character in desert *Drosophila*. Proc Natl Acad Sci U S A 98, 8692–8696 (2001).

29. Karlsson, C. et al. Proteomic Analysis of the *Drosophila* Larval Hemolymph Clot. Journal of Biological Chemistry 279, 52033–52041 (2004).

30. Dziedziech, A., Shivankar, S. & Theopold, U. *Drosophila* melanogaster Responses against Entomopathogenic Nematodes: Focus on Hemolymph Clots. Insects 2020, Vol. 11, Page 62 11, 62 (2020).

31. Lagrange, J., Lecompte, T., Knopp, T., Lacolley, P. & Regnault, V. Alpha-2-macroglobulin in hemostasis and thrombosis: An underestimated old double-edged sword. Journal of Thrombosis and Haemostasis 20, 806–815 (2022).

32. Pike, R. N., Buckle, A. M., le Bonniec, B. F. & Church, F. C. Control of the coagulation system by serpins. FEBS J 272, 4842–4851 (2005).

33. Theopold, U., Li, D., Fabbri, M., Scherfer, C. & Schmidt, O. The coagulation of insect hemolymph. Cellular and Molecular Life Sciences 59, 363–372 (2002).

34. Furie, B. & Furie, B. C. The Molecular Basis of Blood Coagulation Review. Cell 53, 505–518 (1988).

35. Kattula, S., Byrnes, J. R. & Wolberg, A. S. Fibrinogen and Fibrin in Hemostasis and Thrombosis. Arteriosclerosis, Thrombosis, and Vascular Biology vol. 37 e13–e21 Preprint at 10.1161/ATVBAHA.117.308564 (2017).

36. Hanington, P. C. & Zhang, S.-M. The Primary Role of Fibrinogen-Related Proteins in Invertebrates Is Defense, Not Coagulation. J Innate Immun 3, 17–27 (2011).

37. Bono, J. M., Matzkin, L. M., Hoang, K. & Brandsmeier, L. Molecular evolution of candidate genes involved in post-mating-prezygotic reproductive isolation. J Evol Biol 28, 403–414 (2015).

38. Ravi Ram, K., Ji, S. & Wolfner, M. F. Fates and targets of male accessory gland proteins in mated female *Drosophila* melanogaster. Insect Biochem Mol Biol 35, 1059–1071 (2005).

39. Wolfner, M. F. Precious Essences: Female Secretions Promote Sperm Storage in *Drosophila*. PLoS Biol 9, e1001191 (2011).

40. McCullough, E. L. et al. The life history of *Drosophila* sperm involves molecular continuity between male and female reproductive tracts. Proc Natl Acad Sci U S A 119, (2022).

41. Corrigan, L. et al. BMP-regulated exosomes from *Drosophila* male reproductive glands reprogram female behavior. Journal of Cell Biology 206, 671–688 (2014).

42. Hopkins, B. R. et al. BMP signaling inhibition in *Drosophila* secondary cells remodels the seminal proteome and self and rival ejaculate functions. Proc Natl Acad Sci U S A 116, 24719–24728 (2019).

43. Delbare, S. Y. N., Jain, A. M., Clark, A. G. & Wolfner, M. F. Transcriptional programs are activated and microRNAs are repressed within minutes after mating in the *Drosophila melanogaster* female reproductive tract. BMC Genomics 2023 24:1 24, 1–15 (2023).

44. McDonough-Goldstein, C. E., Pitnick, S. & Dorus, S. *Drosophila* female reproductive glands contribute to mating plug composition and the timing of sperm ejection. Proceedings of the Royal Society B: Biological Sciences 289, (2022).

45. Schneider, M. R., Mangels, R. & Dean, M. D. The molecular basis and reproductive function(s) of copulatory plugs. Mol Reprod Dev 83, 755–767 (2016).

46. Avila, F. W., Wong, A., Sitnik, J. L. & Wolfner, M. F. Don’t pull the plug! The *Drosophila* mating plug preserves fertility. Fly (Austin) 9, 62 (2015).

47. Meslin, C. et al. Structural complexity and molecular heterogeneity of a butterfly ejaculate reflect a complex history of selection. Proc Natl Acad Sci U S A 114, E5406–E5413 (2017).

48. Chapin, J. C. & Hajjar, K. A. Fibrinolysis and the control of blood coagulation. Blood Rev 29, 17 (2015).

49. Kelleher, E. S. & Markow, T. A. Reproductive Tract Interactions Contribute to Isolation in *Drosophila*. Fly (Austin) 1, 33–37 (2007).

50. Dorus, S., Evans, P. D., Wyckoff, G. J., Sun, S. C. & Lahn, B. T. Rate of molecular evolution of the seminal protein gene SEMG2 correlates with levels of female promiscuity. Nature Genetics 2004 36:12 36, 1326–1329 (2004).

51. Xu, P. et al. Stable isotope labeling with amino acids in *Drosophila* for quantifying proteins and modifications. J Proteome Res 11, 4403–4412 (2012).

52. Schober, F. A. et al. Stable Isotope Labeling of Amino Acids in Flies (SILAF) Reveals Differential Phosphorylation of Mitochondrial Proteins Upon Loss of OXPHOS Subunits. Molecular & Cellular Proteomics 20, 100065 (2021).

53. Schober, F. A., Atanassov, I., Freyer, C. & Wredenberg, A. Quantitative Proteomics in Drosophila with Holidic Stable-Isotope Labeling of Amino Acids in Fruit Flies (SILAF). in Methods in Molecular Biology vol. 2192 75–87 (Humana Press Inc., 2021).

54. Chang, Y. C. et al. Evaluation of *Drosophila* metabolic labeling strategies for in vivo quantitative proteomic analyses with applications to early pupa formation and amino acid starvation. J Proteome Res 12, 2138–2150 (2013).

55. Coleman, J. M., Benowitz, K. M., Jost, A. G. & Matzkin, L. M. Behavioral evolution accompanying host shifts in cactophilic *Drosophila* larvae. Ecol Evol 8, 6921–6931 (2018).

56. Hughes, C. S. et al. Ultrasensitive proteome analysis using paramagnetic bead technology. Mol Syst Biol 10, 757 (2014).

57. Benowitz, K. M., et al. Chromosome-length genome assemblies of cactophilic *Drosophila* illuminate BioRxiv (2022) doi:10.1101/2022.10.16.512445.

58. Perez-Riverol, Y. et al. The PRIDE database resources in 2022: a hub for mass spectrometry-based proteomics evidences. Nucleic Acids Res 50, D543–D552 (2022).

59. Bolger, A. M., Lohse, M. & Usadel, B. Trimmomatic: A flexible trimmer for Illumina sequence data. Bioinformatics 30, 2114–2120 (2014).

60. Li, H. & Durbin, R. Fast and accurate short read alignment with Burrows– Wheeler transform. Bioinformatics 25, 1754–1760 (2009).

61. Broad Institute. Picard Toolkit. Preprint at (2019).

62. Choi, S., Kim, H. & Paek, E. ACTG: novel peptide mapping onto gene models. Bioinformatics 33, 1218–1220 (2017).

63. Dainat, J. Another Gff Analysis Toolkit to handle annotations in any GTF/GFF format. (Version v0.6.0). Zenodo.

64. Quinlan, A. R. & Hall, I. M. BEDTools: a flexible suite of utilities for comparing genomic features. Bioinformatics 26, 841–842 (2010).

65. Li, H. et al. The Sequence Alignment/Map format and SAMtools. Bioinformatics 25, 2078–2079 (2009).

66. Koboldt, D. C. et al. VarScan 2: Somatic mutation and copy number alteration discovery in cancer by exome sequencing. Genome Res 22, 568– 576 (2012).

67. Cingolani, P. et al. A program for annotating and predicting the effects of single nucleotide polymorphisms, SnpEff: SNPs in the genome of *Drosophila* melanogaster strain w1118; iso-2; iso-3. Fly (Austin) 6, 80 (2012).

68. Kim, D., Paggi, J. M., Park, C., Bennett, C. & Salzberg, S. L. Graph-based genome alignment and genotyping with HISAT2 and HISAT-genotype. Nature Biotechnology 2019 37:8 37, 907–915 (2019).

69. Diaz, F. et al. Divergent evolutionary trajectories shape the postmating transcriptional profiles of conspecifically and heterospecifically mated cactophilic *Drosophila* females. Communications Biology 2022 5:1 5, 1–13 (2022).

70. Bates, D., Mächler, M., Bolker, B. & Walker, S. Fitting Linear Mixed-Effects Models using lme4. J Stat Softw 67, 1–48 (2015).

71. Fox, J. & Weisberg, S. An R Companion to Applied Regression. (2019).

72. Hothorn, T., Bretz, F. & Westfall, P. Simultaneous inference in general parametric models. Biom J 50, 346–363 (2008).

73. Bindea, G. et al. ClueGO: a Cytoscape plug-in to decipher functionally grouped gene ontology and pathway annotation networks. Bioinformatics 25, 1091 (2009).

74. Jones, P. et al. InterProScan 5: genome-scale protein function classification. Bioinformatics 30, 1236–1240 (2014).

75. Liao, Y., Wang, J., Jaehnig, E. J., Shi, Z. & Zhang, B. WebGestalt 2019: gene set analysis toolkit with revamped UIs and APIs. Nucleic Acids Res 47, W199–W205 (2019).

76. Bassett, A. R., Tibbit, C., Ponting, C. P. & Liu, J. L. Highly Efficient Targeted Mutagenesis of *Drosophila* with the CRISPR/Cas9 System. Cell Rep 4, 220–228 (2013).

77. Schneider, C. A., Rasband, W. S. & Eliceiri, K. W. NIH Image to ImageJ: 25 years of image analysis. (2012) doi:10.1038/nmeth.2089.

78. Henrik, S., Bolker, B., Westfall, J. & Aust, F. afex: Analysis of factorial experiments. Preprint at (2016).

79. Lenth, R. Emmeans: estimated marginal means. Aka Least-Squares Means. https://cran.r-project.org/package=emmeans (2019).

80. Brooks, M. E. et al. glmmTMB Balances speed and flexibility among packages for zero-inflated generalized linear mixed modeling. R J 9, 378– 400 (2017).

81. Markow, T. A., Beall, S. & Matzkin, L. M. Egg size, embryonic development time and ovoviviparity in *Drosophila* species. J Evol Biol 22, 430–434 (2009).

