## Supplementary information for "Females translate male mRNA transferred during mating"

**This PDF file includes:**

Supplementary Text
Figures S1 to S4
Tables S1 to S4
Legends for Supplementary files 1 to 4

**Supplementary text**
**Analysis pipelines**

*Filtering for polymorphism at diagnostic peptides*

```
#Trimming reads with Trimmomatic
java -jar trimmomatic-0.39.jar PE -threads 8 ANZA_THX_CKDL210002758-1a_HVCVFDSXY_L4_1.fq.gz ANZA_THX_CKDL210002758-1a_HVCVFDSXY_L4_2.fq.gz ANZA_THX_CKDL210002758-1a_HVCVFDSXY_L4_1.fp.gz ANZA_THX_CKDL210002758-1a_HVCVFDSXY_L4_1.fu.gz ANZA_THX_CKDL210002758-1a_HVCVFDSXY_L4_2.rp.gz ANZA_THX_CKDL210002758-1a_HVCVFDSXY_L4_2.ru.gz ILLUMINACLIP:../jbrowsemapping/adapters/TruSeq3-PE-2.fa:2:30:10:2:keepBothReads LEADING:3 TRAILING:3 MINLEN:36

#Mapping trimmed reads with bwa
bwa mem -t 24 /bwaindexes/ARI ANZA_THX_CKDL210002758-1a_HVCVFDSXY_L4_1.fq.fp.gz ARI ANZA_THX_CKDL210002758-1a_HVCVFDSXY_L4_2.fq.fp.gz | samtools sort -m 2G -@ 4 -o MOV_THX_ONTOARI.bam -

#Removing PCR duplicates using Picard
java -jar /picard.jar MarkDuplicates I= MOV_THX_ONTOARI.bam O= MOV_THX_ONTOARI.nodup.bam M= MOV_THX_ONTOARI.nodup_metrics.txt REMOVE_DUPLICATES=TRUE

# Converting gff to gtf using AGAT
singularity exec /AGAT06.sif agat_convert_sp_gff2gtf.pl -gff /ARI93_cat5_minimal.gff -o /ARI93_cat5_minimal.AGAT.gtf

#Running ACTG to map peptides
java -Xmx6G -Xss2m -jar /ACTG_construction.jar const_params.xml
java -Xmx6G -Xss2m -jar /ACTG_mapping.jar mapping_params.xml
```

const_params.xml file

```
<?xml version="1.0" encoding="UTF-8"?>
<Params>
	<Construction>
		<Inputs>
			<!-- Path of a folder which contains transcriptome models. -->
			<Input format="GTF" type="transcriptome">/ACTG/GTFARI</Input>
			<!-- Path of a folder which contains reference genome. -->
			<Input format="FASTA" type="referenceGenome">/ACTG/FASTAARI/</Input>
		</Inputs>

		<Outputs>
			<!-- Name of a constructed variant splice graph file. -->
			<Output format="ser" type="graphFile">/ACTG/MOVARIpep.ser</Output>
		</Outputs>
	</Construction>

	<Systems>
		<!-- Setting the number of threads. A proper setting can increase the speed of construction. -->
		<TheNumberOfThreads>28</TheNumberOfThreads>
	</Systems>
</Params>
```

mapping_params.xml file

```
<?xml version="1.0" encoding="UTF-8"?>
```

```
<Params>
	<Mapping>
		<Environment>
			<!--
			PV: Mapping [P]rotein database first, then next [V]ariant splice graph.
			PS: Mapping [P]rotein database first, then next [S]ix-frame translation.
			VO: Mapping [V]ariant splice graph [O]nly.
			SO: Mapping [S]ix-frame translation [O]nly.
			-->
			<MappingMethod>VO</MappingMethod>

			<!-- Yes/No: consider isoleucine and leucine as the same. -->
			<ILSame>no</ILSame>

			<!-- Path of a peptide list file that a user wants to map. -->
			<Input format="list" type="peptideList">/ACTG/peptideseq.txt</Input>

			<!-- Path of a folder where the output files will be located. -->
			<Output type="outputPath">/outputMOVARI/</Output>
		</Environment>

		<ProteinDB>
			<!-- If a user sets "MappingMethod" as P[V|S], then the user should provide a folder path containing protein database. -->

			<Input format="FASTA" type="proteinDB">/ACTG/ProteinDBARI/</Input>

			<!-- Yes/No: automatically consider single amino-acid variant when mapping on protein database. -->
			<SAVs>yes</SAVs>
		</ProteinDB>

		<VariantSpliceGraph>
			<!-- Serialization file path of the variant splice graph. -->
			<Input format="ser" type="graphFile">/ACTG/MOVARI.ser</Input>

			<!-- Yes/No: consider junction variation events. -->
			<JunctionVariation>yes</JunctionVariation>

			<!-- Yes/No: consider single exon skipping events. -->
			<ExonSkipping>yes</ExonSkipping>

			<!-- Yes/No: consider intron mapping (exon-extension) events. -->
			<IntronMapping>yes</IntronMapping>

			<!-- If a user gives a VCF (variant call format), then this setting must be "yes". -->
			<Mutation>no</Mutation>

			<!-- If "Mutation" tag is set as "yes", then a user should provide the path of folder where VCF files are. -->
			<Input format="VCF" type="mutation">/ACTG/VCFARI</Input>
		</VariantSpliceGraph>

		<SixFrameTranslation>
			<!-- If a user sets "MappingMethod" as PS or SO, then the user should provide the path of folder where reference genome files are. -->
			<Input format="FASTA" type="referenceGenome">/ACTG/FASTAARI</Input>
		</SixFrameTranslation>
	</Mapping>

	<Systems>
		<!-- A user can set the number of threads. A proper setting can increase the speed of mapping. -->
		<TheNumberOfThreads>28</TheNumberOfThreads>
	</Systems>
</Params>
```

```
#Extracting candidate gene regions from genome using bedtools2
intersectBed -a MOV_THX_ONTOARI.nodup.bam -b peptidelist.bed > MOV_THX_ONTOARI.nodup.candidates.bam
```

```
#mpileup generation using samtools
samtools mpileup -B -Q 0 -f ARIdenovo_r0.93.fasta
MOV_THX_ONTOARI.nodup.candidates.bam >
MOV_THX_ONTOARI.nodup.candidates.mpileup

#Variant calling using VarScan2
java -jar VarScan.v2.4.4.jar mpileup2snp MOV_THX_ONTOARI.nodup.candidates.mpileup --
min-coverage 1 --min-var-freq 0.0001 --p-value 1.0 --output-vcf 1 >
MOV_THX_ONTOARI.nodup.candidates.vcf

#Add the ACTG peptide ID to the vcf file using vcftools
singularity run vcftools.sif
cat /MOV_THX_ONTOARI.nodup.candidates.vcf | vcf-annotate -a /peptidelist.sort.bed.gz \
  -d key=INFO,ID=ANN,Number=1,Type=Integer,Description='ACTG Peptide ID' \
  -c CHROM,FROM,TO,INFO/ANN > /MOV_THX_ONTOARI.nodup.candidates.ID.vcf

#Using SnpEff to identify synonymous and nonsynonymous changes
java -Xmx10g -jar /snpEff/snpEff.jar ARI /MOV_THX_ONTOARI.nodup.candidates.ID.vcf >
/MOV_THX_ONTOARI.nodup.candidates.ID.ann.vcf
```

*RNA-seq of mated female reproductive tracts*

```
#Mapping of RNA-seq reads using HISAT2
cat filelist2.txt | parallel -N 4 -j1 "hisat2 -p 24 -x /Genomes/hisatindexes/{1} -1 {2} -2 {3} --fr
--rna-strandness FR | samtools sort -m 2G -@ 4 -o {4} -"
```

filelist2.txt

```
ARI
MmAs_45_T1_1.fq.fp.gz
MmAs_45_T1_2.fq.rp.gz
MOVARI_45_R1.bam
ARI
MmAs_45_T2_1.fq.fp.gz
MmAs_45_T2_2.fq.rp.gz
MOVARI_45_R2.bam
ARI
MmAs_45_T3_1.fq.fp.gz
MmAs_45_T3_2.fq.rp.gz
MOVARI_45_R3.bam
ARI
MmAs6hT1_CKDL190140453-1a_H3V7FBBXX_L4_1.fq.fp.gz
MmAs6hT1_CKDL190140453-1a_H3V7FBBXX_L4_2.fq.rp.gz
MOVARI_6_R1.bam
ARI
MmAs6hT2_CKDL190140454-1a_H3V7FBBXX_L4_1.fq.fp.gz
MmAs6hT2_CKDL190140454-1a_H3V7FBBXX_L4_2.fq.rp.gz
MOVARI_6_R2.bam
ARI
MmAs6hT3_CKDL190140455-1a_H3V7FBBXX_L4_1.fq.fp.gz
MmAs6hT3_CKDL190140455-1a_H3V7FBBXX_L4_2.fq.rp.gz
```

MOVARI_6_R3.bam

```
#Removal of PCR duplicates using Picard
java -jar /picard.jar MarkDuplicates I=/ARI2020F.bam O=/ARI2020F.nodup.bam
M=ARI2020F.nodup.txt REMOVE_DUPLICATES=TRUE

#Merge bam files
samtools merge /ARI2020F_ARIwithinline.nodup.bam /ARI2020F.nodup.bam
/ARIwithinline_final_dedup.bam

#Exporting exon regions from pooled with D. arizonae bam
intersectBed -a /ARI2020F_ARIwithinline.nodup.bam -b /ARI93_cat5_minimal.EXONS.bed
>/ARI2020F_ARIwithinline.nodup.EXONS.bam

#Generating mpileup with samtools
samtools mpileup -B -Q 0 -f /ARIdenovo_r0.93.fasta
/ARI2020F_ARIwithinline.nodup.EXONS.bam >
/ARI2020F_ARIwithinline.nodup.EXONS.mpileup

#Generating vcf using VarScan2
java -jar /VarScan.v2.4.4.jar mpileup2snp /ARI2020F_ARIwithinline.nodup.EXONS.mpileup --
min-coverage 1 --min-var-freq 0.0001 --p-value 1.0 --output-vcf 1 >
/ARI2020F_ARIwithinline.nodup.EXONS.vcf

#Exporting exon regions from head/thorax D. mojavensis mapped onto D. arizonae
intersectBed -a /MOV_THX_ONTOARI.nodup.bam -b /ARI93_cat5_minimal.EXONS.bed >
/MOV_THX_ONTOARI.nodup.EXONS.bam

#Generating mpileup with samtools
samtools mpileup -B -Q 0 -f /Genomes/ARIdenovo_r0.93.fasta
/MOV_THX_ONTOARI.nodup.EXONS.bam > /MOV_THX_ONTOARI.nodup.EXONS.mpileup

#Generating vcf using VarScan2
java -jar /VarScan.v2.4.4.jar mpileup2snp /MOV_THX_ONTOARI.nodup.EXONS.mpileup --min-
coverage 1 --min-var-freq 0.0001 --p-value 1.0 --output-vcf 1 >
/MOV_THX_ONTOARI.nodup.EXONS.vcf

#Calculating REF and ALT frequency using RNAseq data
perl -ne '@a=split " ";$REFREAD=$a[4]=~tr/.,//;  ;$A=$a[4]=~tr/Aa//; ;$T=$a[4]=~tr/Tt//;
;$G=$a[4]=~tr/Gg//; ;$C=$a[4]=~tr/Cc//; ;$F= $REFREAD / $a[3]; ;$ALTREAD = $A + $T + $G
+$C; print "$a[0] $a[1] $a[2] $a[3] $REFREAD $F $ALTREAD $A $T $G $C\n"' INPUT.mpileup
> OUTPUT.txt
#Note: If the last seven columns are zeros that means that there is an indel or some other issue
and site was not used. The REF_FREQ is calculated using total depth not excluding CIGAR “N.”
Subsequently generated a new column called ADJ_REF_FREQ, which will be equal to
REF_READ/(REF_READ + ALT_READ). Additionally, added an ADJ_Depth column, which is
equal to REF_READ + ALT_READ
```

*Protein domain enrichment analyses*

```
#InterProScan of D. arizonae
```

```
./interproscan.sh -i ARIProteinTRandGN.fasta --goterms -f tsv --cpu 24 -o ARIinterpro5-59-
91.tsv

#R command line for WebGestaltR
library(WebGestaltR)
WebGestaltR(enrichMethod="ORA", organism="others", enrichDatabaseFile =
"/Documents/webGestalR/ARIinterpro_plus_mdFTP_SPLIT_trans.gmt",
enrichDatabaseDescriptionFile = "/Documents/webGestalR/description.des", interestGeneFile =
"/Documents/webGestalR/genelist_mdFTP_all.txt", referenceGeneFile =
"/Documents/webGestalR/reference.txt")
```

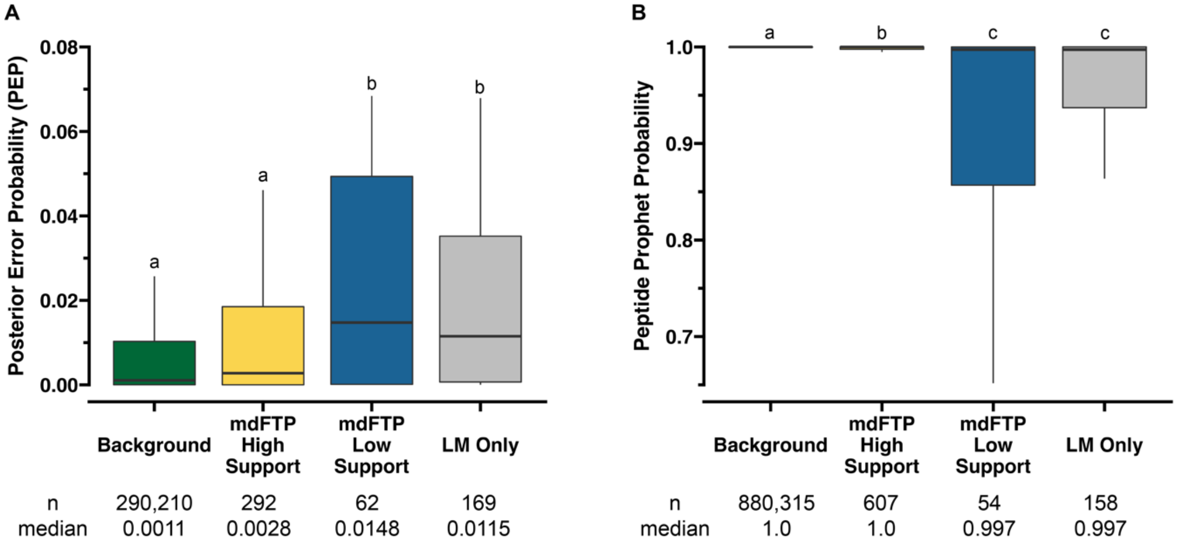


**Figure S1. mdFTP PSMs were of higher quality than LM PSMs.** Posterior error probabilities (PEP) from MaxQuant (A) and Peptide Prophet confidence scores (B) for all identified PSMs after FDR correction (0.01 threshold). Background refers to all PSMs that were not associated with mdFTPs or LM. The middle bar is the median and the boxes represent upper and lower quantiles. The whiskers represent 1.5x inter-quartile range. The letters on top of the box indicate significance of pairwise Kruskall-Wallis test (P<0.01). The sample size and median value for each category of values is provided below each graph.

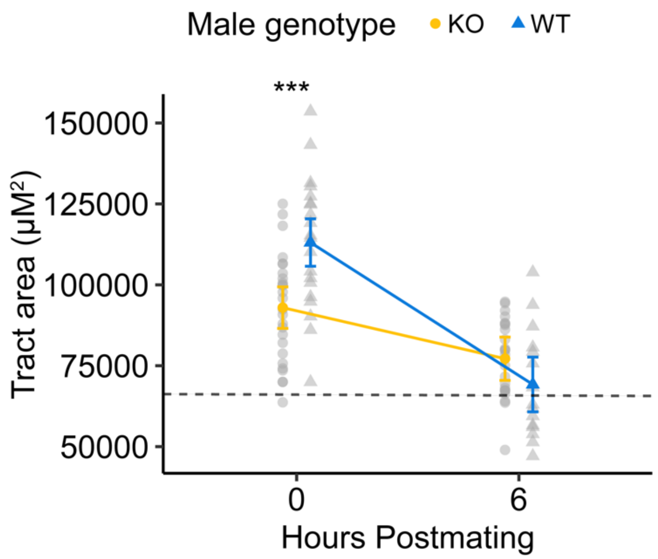


**Figure S2. *ARI/26694* KO effects the size and rate of degradation of the insemination reaction mass.** (a) *Drosophila arizonae* females mated to *D. arizonae* males with an *ARI/26694* KO mutation had a smaller reaction mass initially, as measured by average area, but it degraded more slowly compared to the reaction mass in females mated to WT males (Two-way ANOVA: genotype x time interaction, F= 14.8, *P*= 0.0002). *Post hoc* comparisons were performed using Tukey's method (****P*≤ 0.0001). The dashed line indicates the mean area of unmated female lower reproductive tract. Error bars represent 95% confidence intervals of estimated marginal means. KO-0 hr: n=26; WT-0 hr: n=35; KO-6 hr: n=15

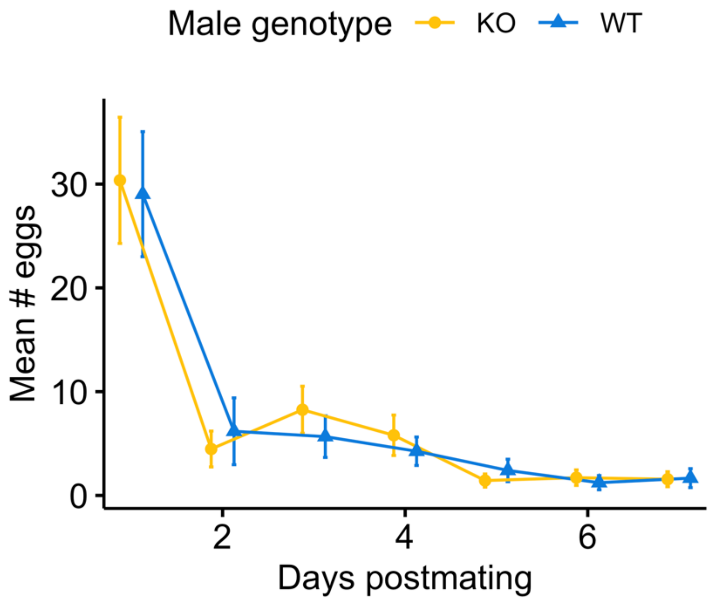


**Figure S3. KO of *ARI/11629* in males did not affect egg-laying by females.** Females mated to KO and WT males produced similar numbers of eggs over seven days postmating (GLMM: Genotype x day interaction, $\chi^2$=0.7, *P*= 0.4145; Genotype, $\chi^2$=1.5, *P*= 0.2193). Error bars represent 95% confidence intervals for the mean. WT: n=52; KO n=69.

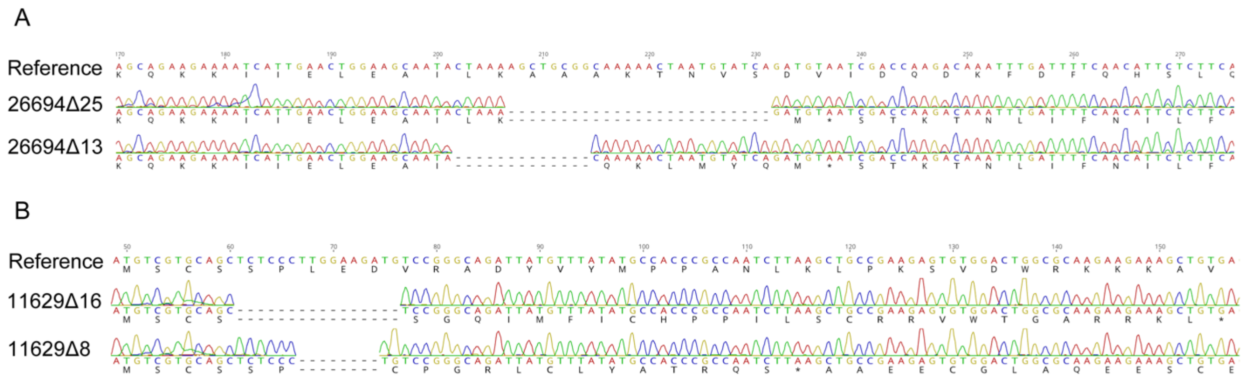


**Figure S4. Mutations in *ARI/26694* and *ARI/11629* generated by CRISPR gene editing.** Two KO mutations were generated for each gene. Flies homozygous for each mutation were crossed to make transheterozgyous males for use in experiments. (A) 25 bp and 13 bp deletion mutations generated in *D. arizonae* (NDSSC: 15081-1271.41) gene *ARI/26694*. Both mutations are within the first ~210 bp of the coding sequence (1341 bp total coding sequence length). Both mutations result in frameshifts that produce a premature stop codon. (B) 16 bp and 8 bp deletion mutations generated in *D. arizonae* (NDSSC: 15081-1271.41) gene *ARI/11629*. Both mutations are within the first ~70 bp of the coding sequence (762 bp total coding sequence length).

| | MaxQuant | | | MSFragger | | |
|---|---|---|---|---|---|---|
| Replicate | 0121 | 0321 | 0222 | 0121 | 0321 | 0222 |
| PSM FDR | 0.01 | 0.01 | 0.01 | 0.01 | 0.01 | 0.01 |
| PSM | 71,891 | 120,553 | 98,286 | 214,368 | 387,875 | 286,891 |
| HA-only PSM | 254 | 372 | 259 | 333 | 575 | 425 |
| LM-only PSM | 31 | 91 | 47 | 39 | 63 | 56 |
| PSM HA-only/LM-only ratio | 8.2 | 4.1 | 5.5 | 8.5 | 9.1 | 7.6 |

**Table S1. Summary of proteomic analysis using multiple pipelines**. Peptide spectrum matches (PSMs) with MS1 intensity quantifications are shown. HA-only refers to PSMs to *D. arizonae* with only heavy label. LM-only refers to PSMs to *D. mojavensis* with only light label.

| | MaxQuant | | | MSFragger | | |
|---|---|---|---|---|---|---|
| Replicate | 0121 | 0321 | 0222 | 0121 | 0321 | 0222 |
| Diagnostic HA peptides | 192 | 249 | 243 | 212 | 196 | 277 |
| Diagnostic HA peptides post-filter | 69 | 67 | 98 | 57 | 61 | 69 |
| Candidate mdFTPs | 64 | 62 | 86 | 46 | 52 | 54 |
| Total unique candidate mdFTPs | | 145 | | | 104 | |
| Total unique candidate mdFTPs post one-hit-wonder filter | | 112 | | | 86 | |

**Table S2. Summary of the filtering of candidate mdFTPs for both analysis pipelines**. Diagnostic HA peptides match the *D. arizonae* (male) peptide sequence but carry the heavy (female) isotopic label. Diagnostic HA peptides were filtered to remove peptides that were polymorphic in either species or were differentiated based only on a leucine-isoleucine substitution. Candidate mdFTPs were filtered to remove those identified by a single peptide (i.e. one-hit-wonders).

| mdFTP support | Number of loci | HAM + HA peptides | | | |
| --- | --- | --- | --- | --- | --- |
| | | Mean | Median | Min | Max |
| High support | 67 | 40.5 | 8 | 2 | 534 |
| Low support | 99 | 13.8 | 7 | 2 | 147 |

**Table S3. Summary of total peptide evidence supporting mdFTP identification.** HAM refers to heavy peptides that have the same sequence in *D. arizonae* and *D. mojavensis* (i.e. non-diagnostic heavy peptides). HA refers to diagnostic peptides that match *D. arizonae* sequence (male) but carry the heavy (female) isotopic label.

| Ingredient | Amount |
|---|---|
| Isotopically labeled yeast (lyophilized) | 330 mg |
| Yeast nitrogen base without amino acids | 15 mg |
| Molasses | 470 mg |
| Sterile water | 7.3 ml |
| Tegosept | 10 mg/24.8 μl ethanol |

**Table S4. Recipe for Lys8 food media used to rear larvae.** The recipe makes one vial, which can be used to rear ~100 larvae.

**Supplementary File S1:** Output data from MaxQuant and MSFragger for diagnostic HA PSMs.

**Supplementary File S2:** List of mdFTPs of high and low support identified using MaxQuant and MSFragger. For each analysis pipeline, both separately and combined, the number of peptides observed per gene, number of times found across replicates, and the respective *D. melanogaster* ortholog (if present) is provided.

**Supplementary File S3:** List of identified seminal fluid proteins (SFPs) in *D. arizonae* and their *D. melanogaster* ortholog.

**Supplementary File S4:** Details of the composition of each of the significant functional clusters identified using ClueGo.
